# Contrasting life history strategies explain contrasting phenology of two co-occurring bumble bee species

**DOI:** 10.64898/2026.07.23.739444

**Authors:** Erin D. Treanore, Sylvana Finn, Genevieve Pugesek, Edison Chae, Stuart Farnham, Maria Ostapovich, Elizabeth Crone

**Author notes:** Erin D. Treanore, current address: Doris Duke Foundation, New York NY USA 10022.

## Abstract

Across species, changing environmental conditions are altering major phenological milestones. In seasonal environments, this can involve shifts in the timing of activity onset, growth, reproduction, and senescence; however, predicting these responses can be challenging and often requires species-specific information. In this study, we investigated phenological patterns of bumble bee colony development using observations from wild nests of two common species, *Bombus griseocollis* and *B. impatiens*. Using observations of nest traffic, we documented dates of nest-searching, peak worker activity, first gyne (new queen) production, and colony senescence over three years of field data collection. We also tested whether colony growth was density dependent, because longstanding life history models have shown that colony growth patterns determine the optimal timing of reproduction in constant environments. Here, we present a novel extension of these models, using them to infer that density-independent colony growth should be associated with extended phenology in warmer years, while density-dependent should have consistent phenology. In total, we found 79 wild nests, including 34 reproductive colonies (19 *B. griseocolis* and 15 *B. impatiens*). *Bombus impatiens* colonies were larger, had a longer activity period, and showed density-independent growth and later reproduction in warmer years, while *B. griseocollis* colonies were smaller, shorter-lived, and showed density-dependent growth and similar dates of reproduction across years. Both species had higher gyne production in warmer years. These results highlight the role of life history theory for predicting interspecific variation in phenological changes, as well as a research framework for exploring possible evolutionary mismatches of existing life histories in changing environments.

## Introduction

Changes in the timing (i.e., phenology) of key life history events, such as growth, maturation, and reproduction, are among the most widespread ecological fingerprints of climate change. As the climate warms, seasonal springtime events appear to be shifting, with documented examples including earlier plant flowering and leaf-out (Menzel 2003; Miller-Rushing and Primack 2008), flight onset in insects (Roy and Sparks 2000), and mammal emergence from hibernation (Inouye et al. 2000). Fewer studies have looked for changes in end-of-season activity (Gallinat et al., 2015), but, at least in temperate climates, summer is getting longer, with later documented dates for phenological milestones typically observed in the fall, such as leaf senescence in plants, end of flight period in butterflies and solitary bees, and timing of post-breeding migration in birds (Dorian et al. 2023; Lawrence et al. 2022; Menzel 2003; Zipf et al. 2017). For some species, it is possible that changes in phenology indicate adaptive matching of life histories to new environments (Iler et al. 2021; Michielini et al. 2021). In other species, however, phenological shifts can cause a misalignment between the optimal environmental conditions and the cues that mediate key life history events across developmental stages (Knapp et al. 2022; Lawrence and Soame 2004, Kudo and Ida 2013). Predicting how a species’ phenology will respond to changing environmental conditions is critical for effective conservation strategies but remains challenging for many taxa.

In this paper, we explore empirical patterns of bumble bee (*Bombus*) phenology in relation to interannual environmental variation. To do so, we use observations from wild colonies of two common bumble bee species, *B. impatiens* and *B. griseocollis*, which we found in the field over three sampling years (2019, 2021, and 2023). Bumble bees are a widely studied species of conservation interest due to their economic and ecological importance as pollinators (Velthuis and van Doorn 2006, Goulson et al. 2008) and recently-observed declines of many *Bombus* species (Colla et al. 2012, Kerr et al. 2015). Nonetheless, the data in this paper are unique in the sense of exploring dynamics of naturally established, wild colonies of this social species. Past studies of drivers of abundance and phenology of bumble bees have used population-level surveys of foraging workers (Pawlikowski et al. 2020, Ogilvie and Caradonna 2022, Kudo et al. 2025). These data are much less labor intensive to collect than colony-level studies, but the effective sample size is unclear, since colonies (rather than individual bees) are the unit at which many phenological changes actually occur. Studies of colony-level processes in bumble bees are typically done with lab-reared colonies, derived from either field-caught (e.g., Williams et al. 2012) or commercially-propagated (e.g., Rondeau and Raine 2024) queens. In general, this approach has not been used to study phenology because, by definition, lab-reared colonies are inherently not synchronized with natural phenological markers. Thus, our study introduces novel aspects of the basic biology of bumble bee colony growth, as well as addressing general questions about drivers of phenology.

Across taxa, approaches to studying phenological shifts in changing environments have relied primarily on statistical or physiological models. Statistically-driven models typically evaluate empirical relationships between traits and changes in phenology. For example, one statistical pattern is that warmer conditions can lead to a switch in the number of generations produced per year (voltinism) (Altermatt 2010) and multivoltine insects are also more likely to extend their seasonal activity periods under these warmer conditions (Hassall et al. 2017; Michielini et al. 2021). Bumble bees are primitively eusocial insects in which colonies are founded by queens in spring. These colonies grow through production of several generations of workers before starting to reproduce (i.e., produce males and gynes (new queens)). At the end of the growing season, old queens, males and workers die and only gynes overwinter. If we interpret colonies as multivoltine populations of workers, the implication of this pattern would be that colonies should grow longer and produce more offspring in warmer conditions. Another statistical pattern is that the start of spring activity for aboveground nesting solitary bee species is more sensitive to temperature than belowground solitary nesting species (Dorian et al. 2023; Stemkovski et al. 2020). If we apply this pattern to bumble bee species, it would imply that phenology of above-ground nesting species (such as one of our focal species, *B. griseocollis*) might be more sensitive to weather conditions than below-ground nesting species (such as our other focal species, *B. impatiens*).

Physiologically-driven models are based on known drivers of life history events. For example, in many multivoltine insects (i.e., species that produce several generations within a single year), spring emergence from diapause is cued by temperature (von Schmalensee et al. 2024), but fall entry into diapause is cued at least in part by daylength (Grevstad and Coop 2015; Saunders et al. 2004). For bumble bees, no immediately known studies have directly tested environmental cues’ role in regulating colony development, likely because of the difficulty of doing experiments at the colony level under natural conditions. Instead, studies of lab-reared colonies have tended to focus on endogenous cues that regulate colony life cycle events, such as the ratio of brood to workers and the timing of colony reproduction (Amsalam et al. 2025, Alaux et al. 2005). Population-level studies suggest that cues for spring emergence include snowmelt and soil temperature (Kudo 2014, Kudo and Cooper 2019), although some studies have found relatively little among-year variation in *Bombus* phenology (Pyke et al. 2016, Kudo et al. 2025). Much less is known about cues for end-of-season phenological milestones in bumble bees, but one early study of *B. flavifrons* (Bowers 1985, 1986) found that colonies in resource-rich environments reproduced earlier in the year, but did not produce more offspring, compared to colonies in resource-poor environments. Another recent study (Riaño-Jiménez et al. 2020) showed earlier reproduction of *B. atratus* in years with more precipitation, which presumbly led to more floral resources. These studies could be interpreted to mean that high floral resources are a physiological cue for earlier reproduction. Such an interpretation would be surprising; intuitively, one might expect that <u>low</u> floral resources would indicate the end of the growing season, and might act as a cue for reproduction. In contrast with these observational studies, one field study with lab-reared colonies of *B. vosnesenskii* showed that early-season resource supplementation caused colonies to produce more queens, with no change in the timing of reproduction (Malfi et al. 2022).

For annual plants and eusocial Hymenoptera, a third approach has evaluated environment-phenology relationships using theoretical models of optimal reproduction. The first of these models, which date back to the 1970s (Cohen 1971; Macevicz and Oster 1976; Oster 1978; Paltridge and Denholm 1974), assumed density independent colony growth, *i.e.,* a constant rate of new worker production per individual worker in a colony, and predicted that the switch to reproduction should be based on cues for the end of the growing season (Amir and Cohen 1990; Mitesser et al. 2007, Verriest et al. 2016). Later extensions of this theory included density dependent colony growth, meaning that colonies experience declining worker recruitment per worker, due to, e.g., limits to queen egg-laying rate (Beekman et al. 1998) or cavity size. In this case, the onset of reproduction is determined by colony size, and is relatively independent of the end of the growing season (Beekman et al. 1998; Hovestadt et al. 2019; Poitrineau et al. 2009).

Based on this body of theory, the presence of density dependence is a critical determinant of bumble bee phenology in relation to changing environmental conditions. For example, rising regional temperatures have been linked to earlier start dates of bumble bee queens activity (Pawlikowski et al. 2020). If the end of activity were determined by density-dependent limits to colony size, these colonies would also switch to reproduction sooner. In a broad sense, this prediction is consistent with studies that found earlier reproduction by *B. flavifrons* and *B. atratus* in higher resource environments (Bowers 1985, 1986, Riaño-Jiménez et al. 2020), if the mechanism of this response were faster colony growth to a critical size. This life history explanation is arguably more intuitively appealing than the idea that high floral resources are a physiological cue for reproduction. In contrast, if the end of activity were determined by the end of the growing season (as predicted with density-independent growth), then the timing of reproduction would not be affected by the start date (Hovestadt et al. 2018). This prediction would be broadly consistent with effects of resource supplementation in *B. vosnesenskii* (Malfi et al. 2022), in which supplemental resources increased colony size and gyne production, but did not affect the timing of reproduction. However, none of these past studies have directly compared relationships between environmental conditions and phenological milestones among species, or related them to mechanisms of colony growth.

In this study, we evaluate patterns of colony growth and phenology using wild nests of two common and co-occurring bumble bee species, *Bombus impatiens* and *B. griseocollis*. We collected detailed demographic data on colony establishment, growth, and reproduction for three years (2019, 2021, and 2023). We estimated timing of colony foundation by surveying springtime queen nest searching for two years (2023 and 2024). We tracked colony growth by monitoring colony traffic (bee entrances and exits) from nest discovery until senescence. To evaluate the mechanisms of colony growth, we tested for evidence of density dependence in colony growth, i.e., declining colony growth rates with increasing colony size, prior to reproduction. We then tested whether the annual timing of worker activity, which we used as a metric for colony size, onset of colony reproduction, and colony senescence differed among years. We predicted that longer colony growth and later reproduction would be associated with longer growing seasons if colony growth was density independent, but not if it was density dependent. Finally, because our study is the first of its kind (in the sense of evaluating growth and reproduction of wild bumble bee colonies in the field), we assessed basic empirical patterns such as the relationship between worker activity and gyne production, and how these differed between species and among years.

## Methods

### Study system

Bumble bees (*Bombus spp.*) inhabiting temperate environments live in annual and simple eusocial colonies that switch from producing workers to sexuals, i.e., males and gynes, before the end of the colony life cycle. In the springtime, queens emerge from winter diapause and search for an appropriate nest site in which to found their colony; location will vary by species but can include locations such as below-ground cavities, trees, or above ground in thatch (Colla et al. 2011). Following nest site selection, a single queen will begin egg laying and engage in nest activities, *e.g*., foraging and brood care, until the first group of female workers emerges. Workers will take over colony tasks while the queen continues laying eggs and the colony grows progressively until the production of sexuals. At this point the gynes will leave the colony to mate and find a suitable hibernaculum to spend the winter months (Alford 1978). Soon after, the colony will senesce, and the annual cycle will repeat the following year.

The two species used in this study, the common eastern bumble bee *(B. impatiens)* and the brown-belted bumble bee (*B. griseocollis)*, are abundant throughout eastern North America (Colla et al. 2011). *Bombus impatiens* forms colonies in nests underground that reach worker populations in the hundreds (Cnaani et al. 2002; Colla et al. 2011). *Bombus griseocollis* colonies are most often found on the surface and thought to be smaller in size (∼50-100 individuals) than *B. impatiens* (Christman et al. 2022; Colla et al. 2011; Plath 1934). In our study system, *Bombus impatiens* nests are found in below-ground cavities (e.g., abandoned rodent burrows) in grasslands and forests, whereas *B. griseocollis* nests are mainly found in thatch in open grasslands (Pugesek and Crone 2021).

Research was conducted on properties of The Trustees of Reservations, a non-profit land trust, located in the immediate region surrounding Ipswich, Massachusetts (USA). Nest monitoring was done in 2019, 2021, and 2023, and queen phenology walks were done in 2023 and 2024. Work was conducted at three separate locations, Appleton Farms (42°38′52.09″ N, 70°51′1.01″ W), Appleton Grass rides (42°38′33.26″ N, 70°51′57.12″ W), and Greenwood Farms (42°41′35.77″ N, 70°48′59.65″ W). Appleton locations were made up of mixed-use agricultural land, natural land cover such as forests, wetlands and meadows, and hay fields for agricultural production. Greenwood Farms is located approximately 10km from the other two sites and has no current land being used for agriculture, instead consisting of deciduous forests, marshes, and grassy meadows.

### Queen phenology walks

To estimate the timing of nest searching behavior for the two species, we performed walks along three standardized transect routes twice a week beginning in mid-April and continuing through the end of spring queen activity (late June) in 2023 and 2024. Routes were located at the three separate locations above, with the routes measuring approximately 3.2 (Appleton Farms), 2.6 (Appleton Farms Grassrides), and 1.3 (Greenwood Farms) kilometers. We walked at a pace of approximately 3 km/hr between 8:00 and 17:30 when temperatures were above 10°C and conditions were dry and calm. We chose this threshold to allow for surveys throughout the nesting season; in our study region, the average daily high temperature is about 12°C in mid-April. During walks, we noted any queen that was flying in an imaginary box that was two meters on either side of the transect and four meters in front of the transect, modeled after the Bumblebee Conservation Trust BeeWalk protocol (Comont and Dickinson 2020). We attempted to net catch every individual encountered, noting the species, and marking them with a bright fluorescent pink powder (DayGlo) to determine if they were recaptured on the same day. As some queens evaded capture or were in a location that was impossible to use a net, *e.g*., a thorn bush, we would take a photo if possible or identify by sight if observed for at least three seconds. We divided queen behavior into stereotypical “nest searching” activity (slow zig-zag flight close to the ground), foraging, and straight-line flight. Because we were interested in the timing of nest establishment (and other behaviors can occur after establishment), we only include queens engaged in stereotypical “nest searching” behavior at the time of observation.

### Nest Identification and Observation

To find nests for observation, we relied on both systematic searches and opportunistic discoveries. Systematic search methods were the same as those conducted by Iles et al. (2019) Pugesek and Crone (2021), and Thuma et al. (2025); these methods were employed across all three study years. In brief, 30 plots 1500m^2^ in size were established evenly across three different habitat types: hay fields, meadows, and forests. Plots were surveyed once a week for four weeks in 2019 and five weeks in 2021 and 2023. In addition to systematic nest searches, free searches of bumble bee nests were also performed (Pugesek and Crone 2021). For further information on nest searching methods see Treanore et al. (2025).

Following confirmation of a nest, nests were observed on a weekly basis between 7:30 and 17:30 when temperatures were above 10°C and conditions were dry and calm. The time at which a nest was observed throughout the day was stratified across the observations to capture differences in activity across the day. Length of each observation was between 20-50 minutes, with <5 out of 1103 total observations falling outside of that window due to shifting weather conditions. During an observation, an observer sat approximately 1-2 meters away from the nest entrance, close enough to observe the individual bee’s caste and behavior, but far enough as to not interfere with behavior. Over the observation period, the observer counted the number of workers, males and gynes entering and exiting. We used these counts as an index of colony size; in cases where colony growth and foraging have been monitored together, worker activity increases with colony size (*e.g.,* Kerr et al. 2019), in spite of the fact that not all workers forage and that we may have seen the same individual workers twice during our monitoring period. Observations ceased when nests senesced, as determined by multiple observation periods in a row with no activity observed. Analyses of colony phenology include only “reproductive” colonies, identified as those at which we observed at least one gyne. We chose to focus on gynes because female-only models are a common first step in life history and demography research (Caswell 2000). As described below, analyses of overall colony gyne production included all observed colonies.

### Data analysis

Statistical analyses and data visualizations were conducted using R statistical software version 4.5.0 (R Core Team, 2025). Across various analyses (described in more detail below), our general approach was to evaluate statistical significance using marginal hypothesis tests, i.e., testing the significance of each factor by comparing the full model to a reduced model with that factor removed. Following standard practice, the full model for each test omitted higher-order terms (i.e., interactions and/or squared terms) involving the factor being tested. Where possible, tests were implemented using the car:Anova function (Fox and Weisberg 2019), which uses this approach as its default test for significance of fixed effects in generalized linear (mixed) models. In other cases (quadratic functions and zero-inflated models, described below), we manually implemented this procedure. In cases where we wanted to compare specific groups, we used *post hoc* comparisons among groups implemented using the emmeans package (Lenth et al. 2019) as a descriptive tool to interpret statistically significant main effects. Code and data used for all analyses are archived on Open Science Framework.

### Phenology

For each species, we estimated the timing of queen nest searching using the biweekly transect walk data and a Generalized Linear Model (GLM) using the glm() function. Day of year (DOY) of each observation (i.e., the day an individual queen was sighted) was set as the response variable, and year, species, site and their interactions, were set as categorical fixed effects. Using data collected during nest observations (see *Methods: Nest Identification and Observations* above), we determined the DOY when worker activity was highest (*Peak worker count*), the first DOY on which a gyne was spotted leaving or entering the natal nest (*First queen observation*), and the approximate DOY of colony senescence, which was defined as > 2 observations with no activity (*Colony senescence date*). These colony-level metrics were each analyzed using a GLM with DOY as the response variable, site as a fixed effect, year as a categorical fixed effect (i.e., comparing differences among years, not a linear trend through time), species as a fixed effect, and an interaction between year and species. We could not include interactions with site for colony-level metrics because colonies of a species were not always present at all sites in all years.

### Colony growth and density dependence

To test for density dependence in colony growth, we isolated the colony growth phase from our weekly nest traffic data. For this analysis, we focused on growth prior to reproduction because neither density-independent nor density-dependent colony growth leads to continued exponential colony growth after the switch to reproduction. Workers take about three and a half weeks to develop from egg to eclosion and have an estimated lifespan of 20-40 days while bumble bee gynes take about five weeks to develop (Brian 1952; Cnaani et al. 2002; Goldblatt and Fell 1987). Therefore, we quantified pre-reproductive colony growth starting from the date on which a nest was found until two weeks prior to the first observed date of gyne activity at each reproductive nest (Fig. 1 A&B). We chose two weeks as an approximate cutoff to capture the final eclosing workers that were laid just prior to gynes; our main conclusions were broadly robust to different choices about the two-week cutoff (in the sense of similar results for choices of 1-3 weeks; Supplemental Table S2.1).

**Figure 1.**
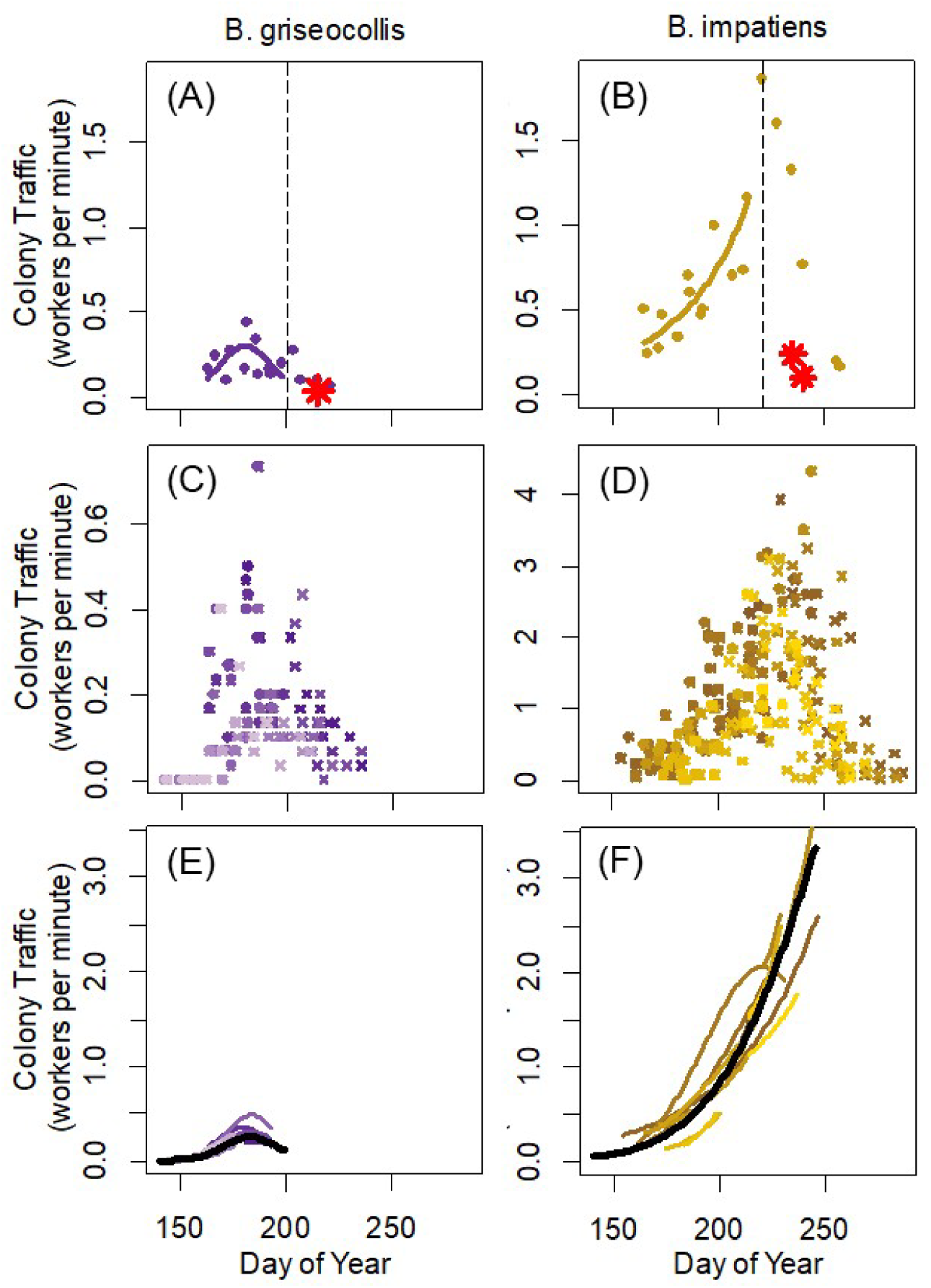
Bumble bee colony traffic. (A&B) Exemplary colonies of (A) *B. griseocollis* and (B) *B. impatiens.* In these panels, red asterisks show queen traffic and filled circles show worker traffic. The dashed line indicates the 2-week cutoff for the “pre-reproductive” period in which we fit colony growth models to data. (C&D) Raw data for all colonies. In these panels, dots are pre-reproductive observations and x’s indicate observations made after the pre-reproductive cutoff. In this row, the two species are shown on different y-axes, so that the shape of seasonal dynamics is visible. (E&F) Pre-reproductive colony growth trajectories predicted from statistical models fit to worker traffic. In both panels, predictions for individual colonies are shown as colored lines, and predictions from fixed effect parameters are shown as solid lines. These predictions are from the full model with quadratic effects, which were strong in *B. griseocollis* and weak in *B. impatiens.* Colors map to the same individual colonies across all panels. Note differences in y-axis values among panels.

Using the colony growth phase data, we modeled colony growth using negative binomial family, log-link generalized linear models (GLMMs; Bates et al. 2009) with worker counts per single observation period for an individual colony as the response variable, and species, DOY, DOY x species, DOY squared and DOY squared x species as predictor variables. Without the squared term, this model predicts exponential colony growth, consistent with predictions from density-independent colony growth models during the pre-reproductive period; in these models, the intercept is the size of a colony (on a log scale) when we first located it, and the slope is the log-scale growth rate per day. The squared term allows this growth rate to slow through time, broadly consistent with density-dependent colony growth. Thus, the test for density dependence is whether the squared term in the model is statistically significant and negative for each species. We chose a quadratic model as a general method to evaluate change in the growth rate that was amenable to being fit to multiple colonies at once using the ‘shrinkage’ properties of GLMMs. Tests for density dependence using the Ricker model, a standard textbook model of density dependence, led to similar conclusions (see *Supplement 1: Additional test for density dependence*).

Our full model included random effects of nest ID, nest ID x DOY, and nest ID x DOY squared, to allow any or all the growth rate parameters to differ among nests. This structure allowed for the initial nest size to be captured by the unique random-effect intercept for each colony and the growth trajectory of each nest to be captured by the unique random effect slopes for each colony (Fig. 1E&F). We also included an offset term (offset(log(duration)) to handle occasional variation in the length of the nest observation periods. We did not include differences among years in this analysis because the total number of colonies for each species was low (see *Results*); therefore, the random effects of nest ID capture both variation among colonies within years and variation among years. Following Zuur et al. (2009), we compared the full model to reduced models with random effects removed; all random effects were supported, so were included in the final data analysis (Supplemental Table S2.2). To illustrate among-colony differences, we extracted the unique predictions for each colony using the predict() function, with the random effect of nest ID set to the value for each colony in sequence (see Fig. 1E&F).

### Worker activity and gyne production

To analyze reproductive performance of each species we used a zero-inflated negative binomial (ZINB) model, implemented with the zeroinfl function from the pscl package in R (Zeileis et al. 2008). We set the total number of gynes sighted at each colony as the response variable, site, species, and year (as a categorical variable) as fixed effects, as well as all two and three-way interactions. Zero-inflated models statistically separate zeroes (in this case colonies where no gynes were sighted) into (1) “true zeros” (colonies that never had gynes) and (2) colonies that produced gynes that we did not observe (Edwards et al. 2021; Zuur et al. 2009). Zero-inflated models estimate two parameters: 1) The proportion of non-reproductive colonies or zeroes and 2) the mean of the non-zero distribution (or the average count of gynes sighted per reproductive colony), around which observations follow a negative binomial distribution that includes some zeros. Post-hoc pairwise comparisons between estimated marginal means were performed using the ‘emmeans’ package (Lenth et al. 2019), using mode = “zero” and mode = “count” to extract the proportion of non-reproductive colonies and average gynes per reproductive colony, respectively.

We also compared the maximum worker activity of colonies that did and did not reproduce using negative binomial GLMs. The response variable in this analysis was the count of workers during the observation with the maximum count of workers per minute (identified in the phenological analysis, above), with an offset of the length of the observation period. These models also included fixed effects of site, species, year (as a categorical predictor), and reproductive status (gynes observed or not), as well as two-and three-way interactions between species, year and reproductive status.

## Results

During our spring phenology walks, we observed a total of 396 nest-searching queens over the two years of observation; 329 of those were *B. impatiens* and 67 were *B. griseocollis.* In total, we found 79 wild bumble bee nests. For *B. griseocollis*, 19 reproductive nests were surveyed a total of 189 times with an average of about 10 times each (median = 9) and the number of observed visits ranging from 1 to 20 per nest (which was largely dependent on when in the season the nest was discovered). Non-reproductive *B. griseocollis* nests (n = 16) were visited a total of 135 times and on average 8.4 times (median = 9.5), with a range of 3 to 12 visits per nest. In comparison, the 15 reproductive *B. impatiens* nests were visited a total of 265 times and an average of 17.7 times each (median = 13) with a range of 5-39 visits per nest. The nonreproductive *B. impatiens* nests were visited a total of 514 times, and on average 17.7 times (median = 21) with a range of 1 to 37 visits per nest.

From these data, analyses of colony phenology included the 34 reproductive colonies (19 *B. griseocollis*; 15 *B. impatiens*). For our analysis of colony growth, we included only reproductive colonies that had sufficient observations prior to colony reproduction. Of the 19 *B. griseocollis* nests that reproduced, 8 were found more than two weeks prior to the first gyne observation and were included in the colony growth analysis. For *B. impatiens,* of the nests that reproduced, 13 were found more than two weeks prior to the first gyne observation and were included in the colony growth analysis. For our analysis of colony size (peak worker activity) and colony reproductive status, we included all 79 colonies (*B. impatiens*: 44, *B. griseocollis*: 35) that were found over the three years of the study. Additional descriptive data for non-reproductive colonies are reported in Supplemental Table S2.3.

### Phenology

The date of *B. impatiens* queen nest searching was slightly later (DOY: 139.81 ± 0.80) than *B. griseocollis* (DOY: 130.22 ± 1.79; (χ^2^□=□9.34, df□=□1, p=0.002, Fig. 2A). The average date of queen nest-searching did not differ between years

**Figure 2.**
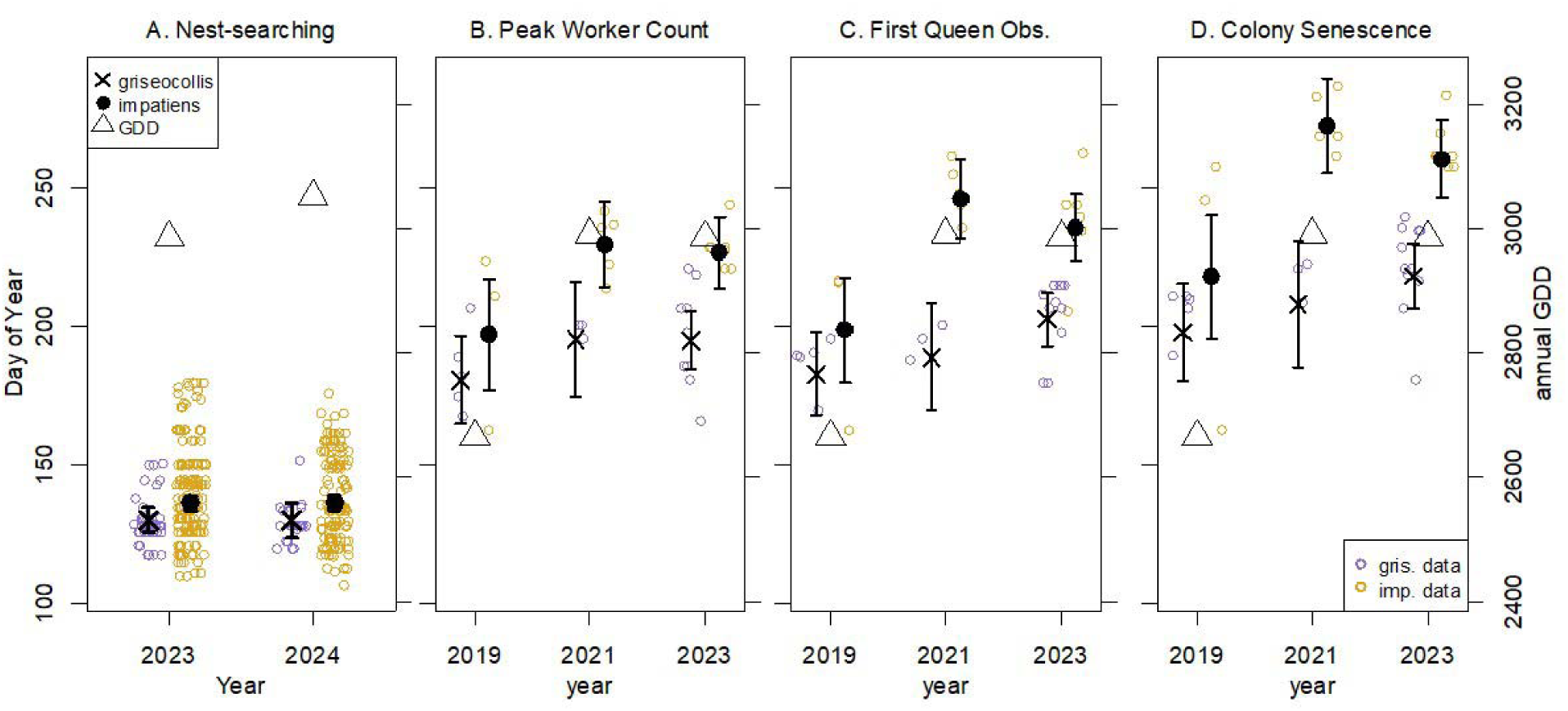
The timing of four key events in the development of *B. griseocollis* and *B. impatiens* reproductive colonies starting with A) Queen nest searching (e.g., colony foundation) B) Peak worker activity C) First gyne observed and D) Colony senescence, as quantified by the date of the last worker observation. Triangles show cumulative annual growing degree days (GDD) scaled to match the secondary y-axis, for comparison with phenology. GDD data were calculated using the Network for Environmental and Weather Applications (NEWA) degree Day Calculator (weather station: Ipswich, Russell Orchards MA USA; newa.cornell.edu) with 4°C as the base temperature. Thick lines show model estimates ± 95% confidence intervals for each species in each year. Colored dots show individual observations (jittered to enhance readability).

(χ^2^□=□0.26, df□=□1, p=0.613), and there was no significant species x year interaction (χ^2^□=0.01, df□=□1, p = 0.926). There was, however, a significant difference between sites (χ^2^□=40.41, df□=□1, p <0.001) and a significant species by site interaction (χ^2^□=8.61, df□=□2, p =0.02), with nest searching by both species occurring earliest at Greenwood Farms, and differing more among sites for *B. impatiens* than *B. griseocollis* (Supplemental Fig. S2.1; Supplemental Table S2.4).

For reproductive colonies, across all life cycle events, the two species differed significantly in their phenology, with *B. griseocollis* occurring earlier in the year; however, the magnitude of this difference was not the same for colony life cycle events (Fig. 2B-D; Supplemental Table S2.5A). *B. griseocollis* colonies had significantly earlier peak worker activity (DOY: 193.84±3.98) compared to *B. impatiens* (DOY: 223.40±4.48, χ^2^□=□25.86, df□=□1, p<0.001, Fig. 2B). Timing of peak worker activity also differed among years (χ^2^□=□9.41, df□=□1, p=0.009), and was earlier in the cooler year (2019) than the other two years (Fig. 2B). There was no species x year interaction (χ^2^□=□1.63, df□=□2, p = 0.484) and no significant difference between sites (χ^2^□=□1.11, df = 2, p = 0.574).

The timing of the first gyne observation differed significantly between species (χ^2^□=□42.21, df□=□1, p<0.001, Fig. 2C), occurring approximately 35 days earlier in *B. griseocollis* colonies (DOY: 198.63 ± 4.52) compared to *B. impatiens* colonies (DOY: 233.87±5.08). Timing of first gyne observation also differed among years (χ^2^□=□16.58, df□=□2, p<0.001), and there was a significant species x year interaction (χ^2^□=□6.99, df□=□2, p=0.03), with *B. impatiens* showing a larger difference in timing between the cooler year (2019) and other two years. The relative timing of first gyne observations of *B. impatiens* was 11.47 ± 14.32 days later than *B. griseocollis* in 2019, 54.00 ± 6.57 days later in 2021, and 33.33 ± 7.17 days later in 2023. Timing of first gyne observations did not differ among sites (χ^2^□=2.53, df□=□2, p=0.28).

Timing of colony senescence differed between species (χ^2^□=□45.29, df□=□1, p<0.001, Fig. 2D), with *B. griseocollis* colonies (DOY: 217.00±5.11) senescing earlier than *B. impatiens* colonies (DOY: 259.53±5.76). Timing of colony senescence also differed among years (χ^2^□=□14.92, df□=□2, p=0.001), but the year x species interaction (χ^2^□=□5.54, df□=□2, p=0.063) and among-site (χ^2^□=□5.04, df□=□2, p=0.080) differences were not statistically significant at the P < 0.05 level. For *B. griseocollis,* the estimated colony activity period (average date of queen nest-searching through average date of senescence) was 87 days, and the average reproductive period (average date of first reproduction through average date of senescence) was 18 days. For *B. impatiens*, the average colony activity period was 119 days, and the average reproductive period was 25 days. Interestingly, these lead to the same amount of overlap of the reproductive period with the whole colony cycle (18/87 = 0.207 and 25/119 = 0.210).

### Colony growth and density dependence

All terms in the colony growth model were retained (Supplemental Table S2.3), based on statistically significant main effects of species (χ^2^□=0□10.05, df□=□1, p = 0.001), species x DOY (χ^2^□=□13.90, df□=□1, p <0.001), and species x DOY^2^ (χ^2^□=□13.50, df□=□1, p < 0.001). *Bombus griseocollis* colony growth rates were significantly density dependent, as evidenced by a significant and negative (slope ± SE =-0.0033±0.0008, Z =-3.949, p <0.001) for the DOY^2^ term (Fig. 1). *Bombus impatiens* colony growth rates were not significantly density dependent (DOY^2^ term: slope ± SE =-0.0002± 0.0002, Z =-1.076, P = 0.282; Fig. 1).

### Worker activity and gyne production

Overall, the number of gynes seen per colony differed between species (χ^2^□=□9.25, df□=□2, p = 0.010) (Fig. 3A&B, Supplemental Table S2.5B), and was higher in *B. impatiens* (2.73 ± 0.76) than *B. griseocollis* (1.79 ± 0.58). This effect primarily reflected a difference in the number of gynes per reproductive nest (marginal test of species for count term only: χ^2^□=□7.38, df□=1, p = 0.007; Fig. 3B), with a weaker difference and species displaying the opposite pattern for the estimated proportion of reproductive nests (marginal test of species for zero term only: χ^2^□=□3.50, df□=□1, p = 0.061; Fig. 3A). Reproduction was somewhat more variable among years in *B. impatiens* than *B. griseocollis*, but this difference was not statistically supported at the P < 0.05 level (species x year interaction: χ^2^ = 6.92, df□=□4, p = 0.140; see Fig. 3A&B). Reproductive output did not differ among sites (χ^2^□=□1.08, df□=□4, p = 0. 898), but was lower in 2019 than 2023 (main effect of year: χ^2^□=□16.01, df□=□4, p = 0.003), primarily due to a lower proportion of reproductive colonies (marginal effect for zero term only: χ^2^□=□8.07, df□=□2, p = 0.018), not a difference in the number of gynes per reproductive colony (marginal effect for count term only: χ^2^□=□2.86, df□=□2, p = 0.239).

**Figure 3.**
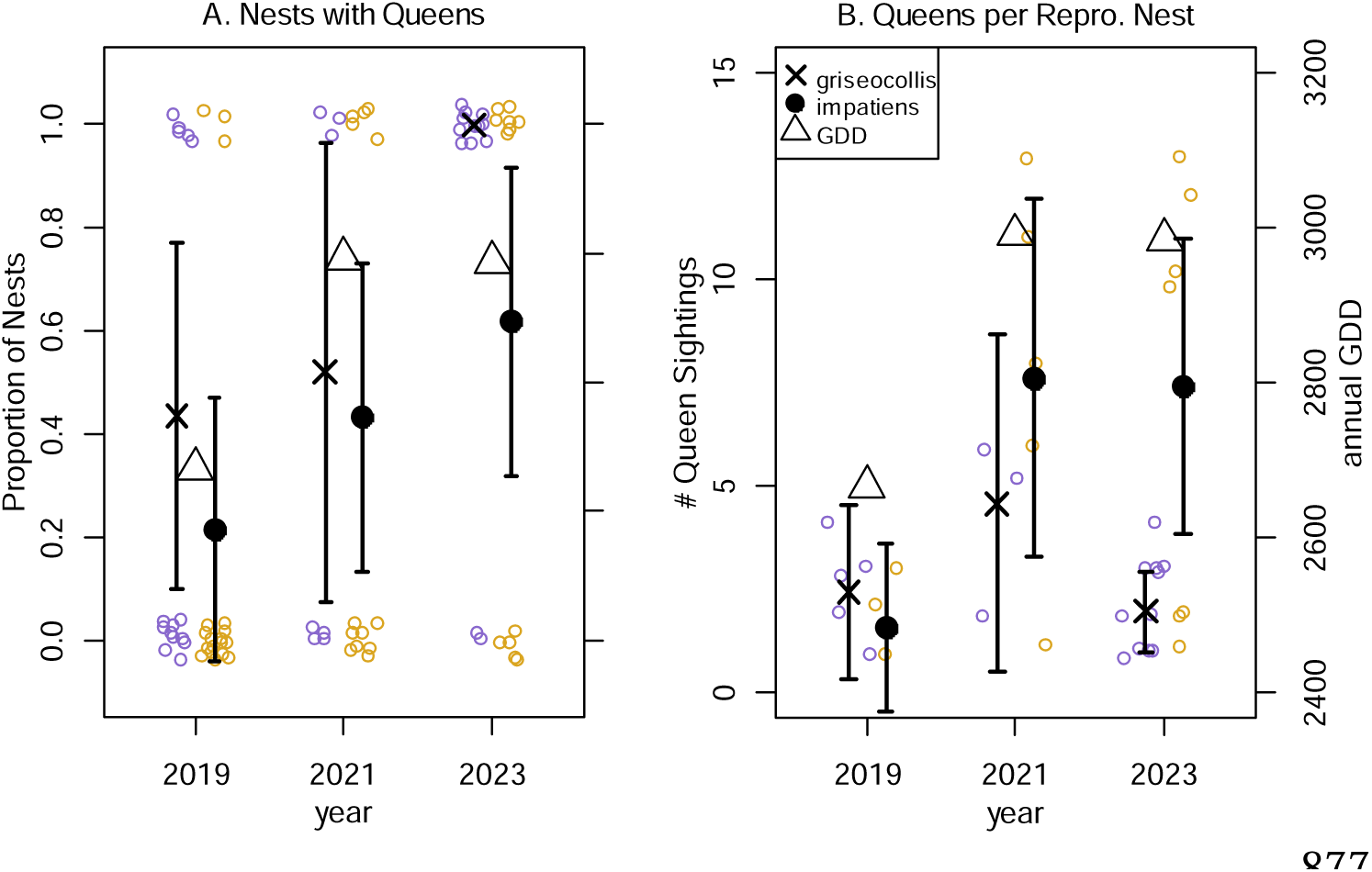
(A) Proportion of colonies that produced queens, and (B) number of queens per reproductive colony, for *B. impatiens* and *B. griseocollis* from the three years of our study. Estimates and 95% confidence intervals are from a zero-inflated negative binomial model, which separates non-reproductive colonies (zero-inflation term) from variation in gyne number among reproductive colonies (count term). Data show observed zeroes and queens sighted. Symbols and colors follow Figure 2. Open circles represent individual nests, jittered for visualization.

Reproductive colonies tended to reach larger sizes than non-reproductive ones (effect of reproductive status on maximum worker count (Fig. 4), although this effect was not statistically significant at the P < 0.05 level (χ^2^□=□3.56, df□=□1, p = 0.059). Maximum worker activity also differed among species (χ^2^□=□111.52, df□=□1, p < 0.001; Fig. 4). There was a tendency toward less difference between reproductive and non-reproductive colonies in 2019 than 2021 or 2023 (year x reproductive status interactions: χ^2^□=□5.04, df□=□2, p = 0.080), especially for *B. impatiens* (Supplemental Fig. 2.2). No other effects were statistically supported predictors of maximum worker activity (see Supplemental Table S2.5C).

**Figure 4.**
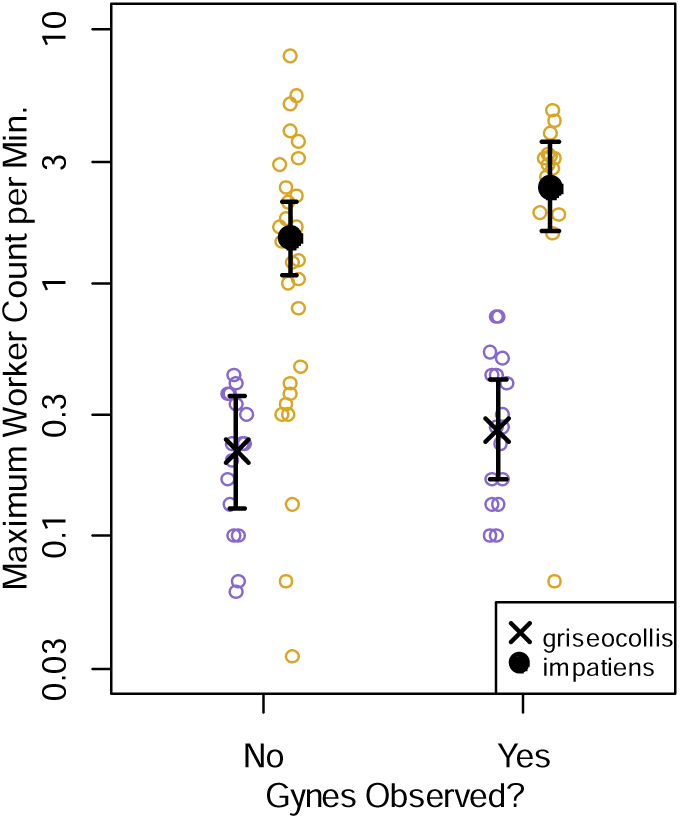
Peak worker foraging rates by reproductive status in *B. impatiens* and *B. griseocollis*. Reproductive colonies (gynes were observed at colony) showed higher peak activity. Open circles show individual colonies; error bars are model-estimated means ± 95% CI from a negative binomial GLM; colors follow Figure 2. Note the log scale on the y-axis.

## Discussion

In our study, *B. impatiens* phenology varied more with growing season length than *B. griseocollis*. Both species reproduced later in warmer years, but this pattern was less pronounced and less statistically supported in *B. griseocollis* than in *B. impatiens*. This difference contrasts with broad expectations from statistical trait-and physiologically-based models of phenology, which in most ways would pool all bumble bees as having similar traits. This difference also occurred in spite of the fact that the two species nest in the same landscape. Therefore, these interspecific differences in phenology were not due to differences in seasonal foraging behavior or elevational differences in climate, in contrast with alpine systems (e.g., Kudo et al. 2025). Of the various approaches that could be used to predict interspecific differences in phenology, the best explanation of our results is expectations from life history models of optimal reproduction. Specifically, *B. impatiens*, the species that extended its colony growth period more in warmer conditions, had density independent colony growth, whereas *B. griseocollis* had density dependent colony growth. Our use of theory in this context is novel; these models were developed primarily to understand the overlap of colony growth and reproduction, and predictions about phenology in relation to the environment are a secondary extrapolation.

Although *B. griseocollis* and *B. impatiens* co-occurred on a common landscape and shared many common traits in our study, one key trait difference, nesting location, might explain their phenological differences, but only in combination with colony growth-reproduction relationships. *Bombus griseocollis* nests mostly in thatch clumps above ground in meadows, whereas *B. impatiens* nests underground in both meadows and adjacent forests. In general, bee species that nest above ground show more phenological variation in relation to the environment (Dorian et al. 2023), which might suggest that *B. impatiens* would be the less responsive species, the opposite of the observed pattern in our study. Similarly, environmental heterogeneity can moderate extreme environmental conditions (Boone et al. 2025), which might also suggest (incorrectly) that since *B. impatiens* nests in both grasslands and forests, it’s phenology would be less variable. However, this difference in nesting location aligns with mechanisms of colony regulation. We typically found *B. griseocollis* colonies in small thatch clumps would not have room for large colonies, whereas *B. impatiens* colonies were often in large underground tunnel systems such as abandoned small mammal burrows, in which colony growth would be less space limited. It is straightforward to hypothesize that these differences in nesting location make *B. griseocollis* colonies more density limited, and that the differences in density dependence allow *B. impatiens* colonies to grow longer when possible.

Our analysis of density dependence relied on worker activity as a metric of colony size. This metric is not perfect; for example, not all workers in a colony contribute actively to foraging (Couvillon and Dornhaus 2010). In spite of this limitation, our results are broadly consistent with past work on these species. A previous study of lab-reared *B. griseocollis* colonies found that colonies never grew larger than 50 individuals (Christman et al. 2022), even when provided with *ad libitum* pollen provisions and artificial nectar, consistent with density dependent colony growth and relatively small colony sizes as inferred from worker traffic in *B. griseocollis*. Lab-reared *B. impatiens* colonies grow to hundreds of workers (e.g., (Cnaani et al. 2002) and are less prone to worker-queen competition (a form of density dependence) than some other species (Cnaani et al. 2002), consistent with our finding of large, density-independent colonies in *B. impatiens*, as inferred from worker traffic. At the same time, use of weekly observations of colony traffic limited our ability to evaluate whether density dependence was associated with graded reproduction (Poitrineau et al. 2009). For *B. griseocollis*, the first gyne observation occurred on a similar day to peak worker activity (Fig. 2 C&D), suggesting simultaneous resource investment in gynes and workers during the preceding weeks. First reproduction occurred later than peak worker activity for *B. impatiens* in the two warmer years (Fig. 2 C&D), which might suggest a discrete switch from growth to reproduction. If the switch to reproduction were more gradual in *B. griseolollis*, we might also expect more overlap of gyne and worker activity (see, *e.g.*, Hovestadt et al. 2019, their Fig. 2), and this was not evident in our data. One possible explanation for this discrepancy is statistical: Dates of first observations are known to be biased later for smaller populations (Edwards and Crone 2022, Primack et al. 2023), and this bias could be extrapolated to smaller bumble bee colonies. More rigorous testing of the overlap of worker and gyne production, which was not the primary aim of our study, would require more intensive monitoring of individual colonies, e.g., daily observations, possibly coupled with mark-recapture observations of individual bees across worker, male and queen castes.

Although we were only able to test some aspects of theory in our system, the idea that different bumble bee species have different life history strategies may inform how we interpret past experimental research. Studies examining factors that regulate the timing of colony development, such as reproduction and the competition point (queen-worker conflict) have investigated control mechanisms such as regulation by the queen, by the workers, or by a colony-level factor, such as ratio of workers to larvae (Alaux et al. 2005; Amsalem et al. 2015). To date, these studies have not reached consensus. A majority of this work, like many other lab-based studies, has been limited to two model species: *B. impatiens* and *B. terrestris* (Treanore et al. 2021). In contrast to *B. impatiens*, *B. terrestris* colony growth patterns mirror what would be expected for density-dependence (Requier et al. 2020). Furthermore, in *B. terrestris,* queen age has also been shown to influence the timing of the switch to reproduction (Alaux et al. 2005); this mechanism could show a similar demographic signal to density dependence. In general, if the two main species used to study reproductive timing have different life history strategies, this could explain why after several decades of work no clear consensus has been reached (Alaux et al. 2005; Beekman et al. 1998).

In addition to timing of reproduction, our study is unusual in that we were able to characterize other features of reproduction for wild bumble bees at the individual nest level. A higher proportion of *B. griseocollis* colonies produced gynes, compared to *B. impatiens*, but fewer gynes were sighted entering/exiting reproductive *B. griseocollis* colonies compared to *B. impatiens* colonies. For both species, a large fraction of colonies did not reproduce, which echoes the limited number of studies that have examined colony reproductive success under natural conditions in temperate environments. For example, a study examining wild bumble bee colonies in the UK found that 21%-71% of colonies produced gynes over a two-year period (Goulson et al. 2018), while another study using artificial domiciles found this range to be between 16%-57% (Richards 1978). We observed that reproductive colonies tended to be larger than non-reproductive colonies but being a larger or more active colony did not always lead to colony reproduction; this pattern was especially noticeable in *B. impatiens*. Other research conducted under similar conditions (Wisconsin; Michigan, USA) but using commercial *B. impatiens* colonies also found high numbers of non-reproductive colonies, and that colony reproduction tended to be associated with larger colonies and greater worker number (Spiesman et al. 2017). Our results and these previous studies demonstrate three important points 1) Not all wild bumble bee colonies reproduce 2) Colony reproductive success varies significantly year-to-year 3) Reproductive colonies tend to be larger. Why some colonies reproduce and others do not is only coarsely understood, but colony nutrient intake quality, quantity, and timing have been frequently cited as playing a role (Hemberger et al. 2020; Lanterman and Goodell 2018; Malfi et al. 2022; Pelletier and McNeil 2003; Vaudo et al. 2018). Our results also suggest that there may be tradeoffs among species in the tendency to produce gynes vs. the number of gynes produced.

Although our study was limited to three years, two of these years were noticeably warmer than the third. Both species demonstrated higher reproductive outputs in warmer years. This result is partly intuitive because the length of the flowering season of many angiosperms, particularly legumes (which are known to produce pollen that is protein-rich and preferentially foraged by bumble bees; Wood et al. 2021), is increasing substantially with warming temperatures (Zhou et al. 2022). Augmented availability of high-quality resources could support colonies of both species in acquiring sufficient energy to support colony reproduction (Cueva del Castillo et al. 2015; Vaudo et al. 2018). In contrast, previous research with bumble bees has generally shown negative effects of warming temperatures on everything from worker physiology to colony foraging behavior (Hemberger et al. 2023; Kuo et al. 2023). One explanation for this disparity is that our research was conducted in in the field, and therefore includes effects on plant quality, as well as direct effects on foragers (Ogilvie et al. 2017). In addition, much of the previous research has focused on extreme temperatures or events, *e.g.*, heatwaves, rather than overall warmer seasonal temperatures. Together, past work and our results may illustrate a “goldilocks” reproductive response, with the conditions observed in our system resulting in increased floral resource availability due to warmer temperatures, even though occasional extreme heat events might tend to have catastrophic effects on individual and colony performance (Thuma et al. 2025).

In closing, our study suggests, first, contrasting responses of *B. impatiens* and *B. griseocollis* to predicted environmental changes. We know that in many regions changing environmental conditions are leading to longer, warmer growing seasons and extended periods of floral resource availability (Ogilvie et al. 2017). Such conditions would likely benefit social bee species that exhibit density-independent growth. Colonies would have a longer period of worker production and could produce a greater number of sexuals, like our observations of *B. impatiens* reproduction. Though not explicitly examined, gyne production and dispersal would theoretically still be synchronized with the end of season conditions that then mediate diapause onset, such as sufficiently cool temperatures (Waybright and Dillon 2024). Conversely, how species with density-dependent patterns of colony growth, like *B. griseocollis*, would respond to warmer, longer, activity periods is not immediately obvious. If colony reproduction occurs earlier in the season, gynes may enter diapause earlier, and possibly even remain in diapause for a longer period. Past research suggests that warmer pre-winter and winter conditions have increased metabolic costs for individual gynes in diapause (Vesterlund et al. 2014) often leading to higher rates of mortality and reduced post-diapause fitness (Nielsen et al. 2022).

In addition, our study suggests new directions for theoretical research. Prior to this study, models of optimal reproduction in social insects have mostly been used to explore the conditions that lead to a sudden vs. gradual shift in allocation of resources from colony growth to reproduction, rather than the timing of reproduction in relation to the environment (but see Hovestadt et al. 2019). Our extrapolation of model predictions to include environmental drivers is heuristic, rather than formally mathematical. It could be especially relevant to adopt an approach similar to evolutionary mismatch theory in behavioral ecology (Pollack et al. 2022), in which models are used evaluate the expected optimal responses to environmental variation in stationary environments, and then to explore the consequences of these strategies for rapidly changing environments (Johansson et al. 2013). We hope our exploration of these issues encourages future research to understand how optimal life cycles in static environments affect expected responses to environmental change.

## Acknowledgements

We would like to acknowledge Krishna Muhundan, Luke Aengenheyster, Jessie Thuma, Kyle Burton, and Heidy Acevedo for assistance in the field. Furthermore, we thank the Trustees of Reservations for their support. Partial funding for this research was provided by Tufts University College of Arts and Sciences and an NSF DEB 2203158 awarded to E. Crone.

## Author Contributions

Elizabeth Crone conceptualized and designed the study; Erin Treanore, Sylvana Finn, Genevieve Pugesek, Edison Chae, Stuart Farnham, and Maria Ostapovich all helped with data collection; Erin D. Treanore and Elizabeth Crone analyzed and interpreted the data; Erin Treanore led the writing of the manuscript. All authors read and approved the final manuscript.

## Data availability statement

Data from this study will be archived and made publicly available at the time of publication on Open Science Framework.

