## Supplemental info for "Contrasting life history strategies explain contrasting phenology of two co-occurring bumble bee species"

**Supplement 2. Supplementary Tables and Figures.**

| Table S2.1. Results of likelihood ratio tests assessing the effects of species identity (species), scaled day of year (scale(doy)), its quadratic term (scale(DOY2)), and their interactions on the number of workers (n_worker) observed across three temporal windows (7, 14, and 21days). Significance codes: ***p < 0.001, **p < 0.01, *p < 0.05, .p < 0.1. | | | | |
| --- | --- | --- | --- | --- |
| Term | **Chisq** | **Df** | **Pr(>Chisq)** | **Significance** |
| 7 days |  |  |  |  |
| species | 29.28 | 1 | 6.27E-08 | *** |
| scale(doy) | 19.53 | 1 | 9.90e-06 | *** |
| scale(DOY2) | 9.42 | 1 | 0.00215 | ** |
| species:scale(doy) | 12.30 | 1 | 0.00045 | *** |
| species:scale(DOY2) | 12.35 | 1 | 0.00044 | *** |
| 14 days |  |  |  |  |
| species | 10.61 | 1 | 0.00113 | ** |
| scale(doy) | 21.39 | 1 | 3.75e-06 | *** |
| scale(DOY2) | 4.08 | 1 | 0.04335 | * |
| species:scale(doy) | 13.90 | 1 | 0.00019 | *** |
| species:scale(DOY2) | 13.50 | 1 | 0.00024 | *** |
| 21 days |  |  |  |  |
| species | 6.19 | 1 | 0.01284 | * |
| scale(doy) | 23.49 | 1 | 1.26e-06 | *** |
| scale(DOY2) | 0.62 | 1 | 0.43243 |  |
| species:scale(doy) | 10.737 | 1 | 0.00105 | ** |
| species:scale(DOY2) | 10.13 | 1 | 0.00146 | ** |

| Table S2.2. Statistics from analysis of colony growth. Sets of random effects were evaluated using the model with all fixed effects. ‘Marginal’ likelihood ratio tests for the fixed effects were calculated using the best combination of random effects. Effects of DOY were evaluated using models with the higher order (DOY^2^) term removed because these are clearly collinear. DOY refers to day of year, and ID refers to unique nest identity. | | | | | | | |
| --- | --- | --- | --- | --- | --- | --- | --- |
| Fixed effects | | | | **Random effects** | | | |
|  | χ^2^ | df | P |  | | ΔAIC* | Np* |
| Species | 10.6 | 1 | 0.001 | None | | 34.1 | 7 |
| DOY | 21.4 | 1 | <0.001 | ID | | 4.2 | 8 |
| DOY^2^ | 4.1 | 1 | 0.043 | ID×DOY | | 2.2 | 10 |
| Species×DOY | 13.9 | 1 | <0.001 | ID×DOY+ID×DOY^2^ | | 0.0 | 13 |
| Species×DOY^2^ | 13.6 | 1 | <0.001 |  |  |  |  |
| * AIC = Akaike’s “An Information Criterion”. Np = Number of estimated parameters | | | | | | | |

| Table S2.3. Descriptive statistics calculated from the raw data for the phenology of colony development in each species, maximum size, and number of queens produced across all colonies, and then exclusively in colonies that reproduced. | | | | |
| --- | --- | --- | --- | --- |
| Colony Metrics | ***Bombus impatiens*** | | ***Bombus griseocollis*** | |
|  | Non-reproductive colonies  *(n=29)* | Reproductive colonies  *(n=15)* | Non-reproductive colonies  *(n=16)* | Reproductive colonies  *(n=19)* |
| Peak Activity (DOY) | 209.03±5.40 | 223.40±4.94 | 188.81±3.21 | 193.84±3.62 |
| First day of reproduction (DOY) | *na* | 233.87±6.56 | *na* | 198.63 ± 3.14 |
| Colony senescence (DOY) | 239.83±6.87 | 259.53±7.47 | 206.06±2.55 | 217.00±3.50 |
| Number of gynesen | *na* | 6.33±1.24 | *na* | 2.63±0.33 |
| Peak colony activity | 1.837±0.234 | 2.835±0.325 | 0.233±0.315 | 0.314±0.289 |

| Table S2.4. Statistics from analysis of queen walks. ‘Marginal’ likelihood ratio tests were implemented with the car::Anova function. | | | |
| --- | --- | --- | --- |
|  | χ^2^ | df | P |
| Site | 40.4 | 2 | <0.001 |
| Species | 9.3 | 1 | 0.002 |
| Year | 0.3 | 1 | 0.613 |
| Site ×Species | 8.1 | 2 | 0.017 |
| Site ×Year | 1.2 | 2 | 0.559 |
| Species×Year | <0.1 | 1 | 0.926 |
| Site ×Species×Year | 0.2 | 2 | 0.902 |

| Table S2.5. Analyses of colony traffic. Phenology metrics were analyzed using reproductive colonies only. Gyne Production and Maximum Worker Count were analyzed for all colonies, in order to compare reproductive and non-reproductive colonies. ‘Marginal’ likelihood ratio tests were implemented using the car::Anova function for Phenology and Maximum Worker Count, and implemented by manually creating nested model pairs for zero-inflated queen production models. | | | | | | | | | |
| --- | --- | --- | --- | --- | --- | --- | --- | --- | --- |
|  | χ^2^ | df | P | χ^2^ | df | P | χ^2^ | df | P |
| A. *Phenology* |  | | |  | | |  | | |
|  | Max. Count DOY | | | First Gyne DOY | | | Senescence DOY | | |
| Site | 1.1 | 2 | 0.574 | 2.5 | 2 | 0.282 | 5.0 | 2 | 0.080 |
| Species | 25.9 | 1 | <0.001 | 42.2 | 1 | <0.001 | 42.3 | 1 | <0.001 |
| Year | 9.4 | 2 | 0.009 | 16.6 | 2 | <0.001 | 14.9 | 2 | 0.001 |
| Species×Year | 1.6 | 2 | 0.484 | 7.0 | 2 | 0.030 | 5.5 | 2 | 0.063 |
| B. *Gyne* C. *Production* | | | | | | | | | |
|  | Overall | | | Zero term | | | Count term | | |
| Site | 1.1 | 4 | 0.898 | 0.5 | 2 | 0.775 | 0.5 | 2 | 0.771 |
| Species | 9.2 | 2 | 0.010 | 3.50 | 1 | 0.061 | 7.4 | 1 | 0.007 |
| Year | 16.1 | 4 | 0.003 | 8.1 | 2 | 0.018 | 2.9 | 2 | 0.239 |
| Species×Year | 6.9 | 4 | 0.140 | 2.5 | 2 | 0.286 | 4.9 | 2 | 0.086 |
| *C. Maximum Worker Count* | | | | | | | | | |
| Site | 2.1 | 2 | 0.348 |  |  |  |  |  |  |
| Species | 111.5 | 1 | <0.001 |  |  |  |  |  |  |
| Year | 2.5 | 2 | 0.284 |  |  |  |  |  |  |
| Reproductive* | 3.6 | 1 | 0.059 |  |  |  |  |  |  |
| Species×Year | 0.1 | 2 | 0.937 |  |  |  |  |  |  |
| Species×Reproductive* | 0.3 | 1 | 0.595 |  |  |  |  |  |  |
| Year×Reproductive | 5.0 | 2 | 0.080 |  |  |  |  |  |  |
| Species×Year×Repro.* | 0.9 | 2 | 0.637 |  |  |  |  |  |  |
| * Reproductive status, classified as colonies at which new gynes were observed vs. colonies at which new gynes were not observed. | | | | | | | | | |

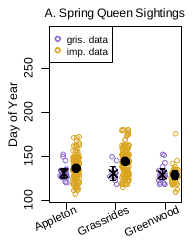

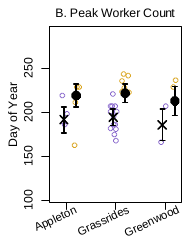

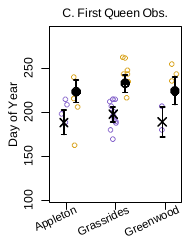

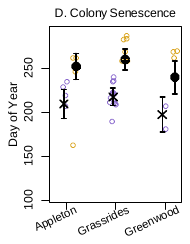

**Figure S2.1.** Differences among sites in timing of four key events in the development of *B. griseocollis* and *B. impatiens* reproductive colonies: A) Queen nest searching (a measure of colony foundation), B) Peak worker activity, C) First gyne (new queen) observed, and D) Last worker observation (a measure of colony senescence). Different sites where nests and queens were observed are listed by name (see main text) on the x-axis.

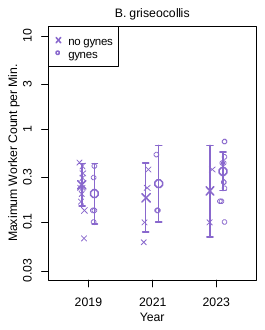

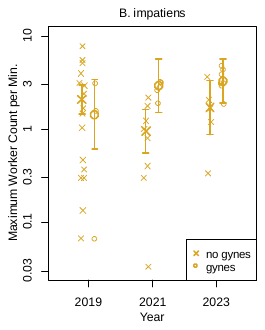

**Figure S2.2.** Peak worker foraging rates by reproductive status in *Bombus impatiens* and *B. griseocollis* broken down by year. Open circles show individual colonies; error bars are model-estimated means ± 95% CI from a negative binomial GLM. Note the log scale on the y-axis.
